# Inferring causality in biological oscillators

**DOI:** 10.1101/2021.03.18.435997

**Authors:** Jonathan Tyler, Daniel Forger, Jae Kyoung Kim

## Abstract

A fundamental goal of biological study is to identify regulatory interactions among components. The recent surge in time-series data collection in biology provides a unique opportunity to infer regulatory networks computationally. However, when the components oscillate, model-free inference methods, while easily implemented, struggle to distinguish periodic synchrony and causality. Alternatively, model-based methods test whether time series are reproducible with a specific model but require inefficient simulations and have limited applicability. Here, we develop an inference method based on a general model of molecular, neuronal, and ecological oscillatory systems that merges the advantages of both model-based and model-free methods, namely accuracy, broad applicability, and usability. Our method successfully infers the positive and negative regulations of various oscillatory networks, including the repressilator and a network of cofactors of pS2 promoter, outperforming popular inference methods. We also provide a computational package, ION (Inferring Oscillatory Networks), that users can easily apply to noisy, oscillatory time series to decipher the mechanisms by which diverse systems generate oscillations.

## Introduction

A fundamental goal in biology is to uncover the causal interactions among system components. To identify the casual interactions, conventional methods require experimental manipulation of one or more components to investigate the effect on others in the system. However, this approach is time-consuming and costly, particularly when the number of components in a system increases. On the other hand, thanks to recent technological advances (e.g., GFP, luciferase, microarray, etc.), measuring time-series data has become relatively easy. Accordingly, inferring direct regulations along with type (positive/negative) solely given time-series data is an important tool to provide key insights into the mechanisms underlying the system in a timely and inexpensive manner (*1*).

The unprecedented growth in the amount of biological data has revealed that biological processes frequently exhibit oscillatory behavior in time-series data, e.g., about half of the protein-coding genome is transcribed rhythmically (*2, 3*). To infer networks from oscillatory data, a popular model-free method, Granger Causality (GC) based on predictability, i.e., *X* causes *Y* if *X* has unique information that can improve the prediction of *Y*, has been used (*4, 5*). However, as GC relies heavily on the assumption that the time-series data are stationary (*6*), it is challenging to apply GC to highly nonstationary oscillatory time-series data (*5, 7–9*). To overcome this limitation of GC, inference methods for dynamical systems, such as Convergent Cross Mapping (CCM), have been developed, based on a differing view of predictability, i.e., *X* causes *Y* if historical values of *X* can be recovered from *Y* alone (*10–20*). Despite the success of CCM methods in many biological applications, they frequently infer interactions between independent components when they oscillate with similar periods due to difficulty in distinguishing synchrony and casual interaction (*21*), indicating that these methods are likely to infer false-positive interactions in oscillatory networks. Nonetheless, these model-free methods remain widely used due to their ease of implementation and broad applicability to a large class of networks.

Alternatively, model-based methods were proposed that infer causality by determining whether time-series data are reproducible with mechanistic models. Testing the reproducibility requires computationally-expensive model simulations and fittings (*22–35*), but, as long as the underlying model is accurate, model-based methods do not suffer from false positive predictions unlike model-free methods. However, the inference results strongly depend on the choice of model, which is frequently based on limited information. Thus, inference methods using more general ODE forms were developed (*36–45*). For example, previously, we developed a method that infers causation from *X* to *Y* by checking whether oscillatory time-series data for *X* and *Y* are reproducible with a common ODE model for biological oscillators: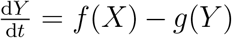, where *f* and *g* describe the synthesis and degradation rates of *Y*, respectively (*41*). Pigolotti et al. (*36*) considered the most general possible mechanistic model between two components:

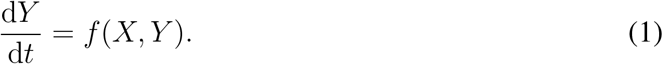

However, unlike (*41*), this method uses only the minima and maxima rather than all of the time-series data (*36*). Thus, the method requires the restrictive assumption that all given components are in a single negative feedback loop (i.e., the method determines the order of given components in a feedback loop). Moreover, extensions of the method (*37, 38*) require that a single negative feedback loop structure drive the dynamics, limiting their applicability.

Here, we develop an inference method for biological oscillators described by Eqn. (1) that merges the advantages of model-based and model-free methods, namely usability, broad applicability, and accuracy, while mitigating the drawbacks of each. Specifically, we identify a fundamental relationship between the general model (Eqn. (1)) and its oscillatory solution. By using this relationship, we develop a simple functional transformation (i.e., regulation-detection function) of a pair of oscillatory time-series data that easily tests whether the time-series data are reproducible with the general model. This transformation enables accurate and precise inference of the (self-)regulation type (e.g., positive, negative, or a mixture) between two components *X* and *Y* described by Eqn. (1). This allows us to infer various network structures such as a cycle, multiple cycles, and a cycle with outputs from *in silico* oscillatory time-series data. Furthermore, our method also successfully infers regulation types from noisy experimental data measured at the molecular and organismal levels. In particular, from time-series data of the repressilator and cofactors at the pS2 promoter, our method infers networks that match current biological knowledge while popular model-free methods incorrectly infer nearly fully connected networks. Importantly, our method predicts hidden regulations for the pS2 promoter after estradiol treatment, which guides investigation. We also provide a user-friendly computational package (ION: Inferring Oscillatory Networks) that implements our method to infer network structures of biological oscillators, which requires minimal user effort.

## Results

### Inferring regulation types from oscillatory time series

In the reduced FitzHugh-Nagumo model (Fig. 1A) (*46*), which describes the interactions between the membrane potential of a neuron (*V*) and the accommodation and refractoriness of the membrane (*W*) (*46, 47*), *W* positively regulates *V* while *V* negatively regulates *W*. In addition, *V* displays a mixture of positive and negative self-regulation while *W* negatively regulates itself.

**Figure 1:**
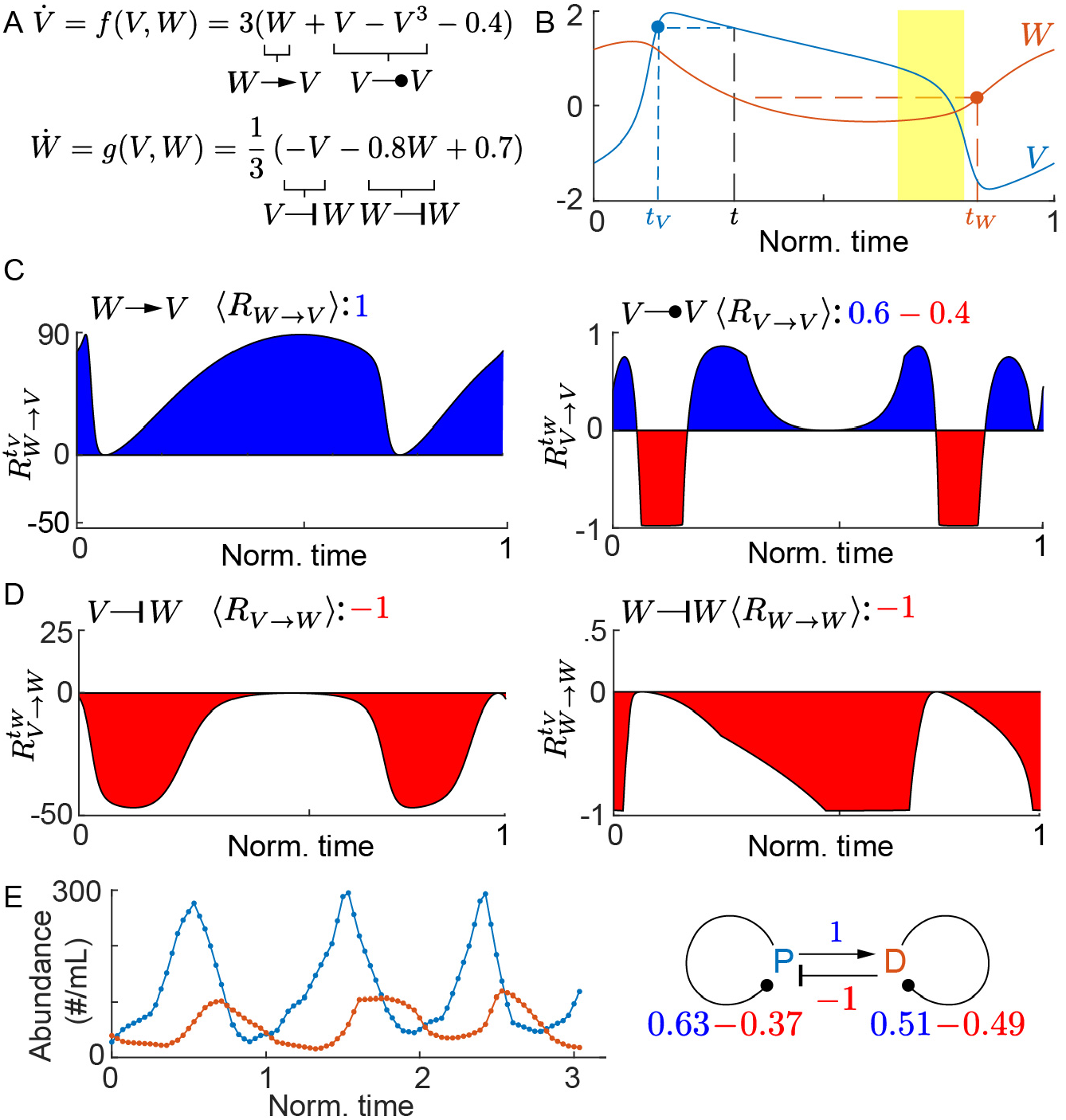
Regulation-detection functions and scores reflect regulation types. (A) The FitzHugh-Nagumo model describes the interactions between the membrane potential of a neuron (*V*) and the accommodation and refractoriness of the membrane (*W*). *W* positively regulates *V* while *V* negatively regulates *W*. In addition, *V* displays a mixture of positive and negative self-regulation while *W* negatively regulates itself. (B) Time series of one cycle simulated with the FitzHugh-Nagumo model. Although *W* positively regulates *V* (i.e., 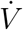 positively depends on *W*), 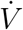 decreases despite increasing *W* (yellow region) because the self-regulation of *V* on 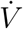 masks the effect of *W* on 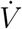. On the other hand, for the the pair of time points *t* and reflection time *t*_*V*_, where *V* (*t*) = *V* (*t*_*V*_), if *W* (*t*) is greater (less) than *W* (*t*_*V*_), then 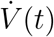 should be greater (less) than 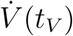. Similarly, as *V* negatively regulates *W*, if *V* (*t*) is greater (less) than *V* (*t*_*W*_), then 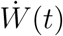 should be less (greater) than 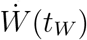 for the pair of time points *t* and *t*_*W*_, where *W* (*t*) = *W* (*t*_*W*_). (C) The regulation-detection function 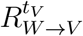 (Eqn. (2)) is positive, and thus the regulation-detection score ⟨*R* _*W*→*V*_ ⟩ (Eqn. (4)) equals one, reflecting the positive regulation of *W* on *V*. The sign of 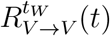 (Eqn. (3)) changes, and thus −1 < ⟨*R*_*V*→*V*_ ⟩ < 1 (Eqn. (4)), reflecting the mixture of positive and negative self-regulation of *V*. (D) Both 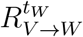 and 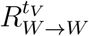 are negative, and thus ⟨*R*_*V*→*W*_⟩ = ⟨*R*_*W*→*W*_⟩ = −1, reflecting the negative regulation of *V* on *W* and the self-regulation of *W*. (E) Regulation-detection scores are calculated from the time-series population data of two bacteria: *Paramecium*, P (blue), and *Didinium*, D (red) (data taken from (*10*)). Reflecting the known predatory interaction, ⟨*R*_*P*→*D*_⟩ = 1 and ⟨*R*_*D*→*P*_ ⟩ = −1. Furthermore, reflecting that the self-regulation of both P and D consists of both positive (i.e., birth) and negative (i.e., death) regulation,⟨*R*_*P*→*P*_ ⟩ = 0.26 and ⟨*R*_*D*→*D*_⟩ = 0.02.

How are such inter- and self-regulations reflected in the oscillatory change of *V* and *W* simulated with the model (Fig. 1B)? The change in *V* and *W* does not directly reflect their regulatory interactions. For instance, although *W* positively regulates *V*, when *W* increases, *V* does not always increase (e.g., in the region highlighted in yellow, Fig. 1B). This is because *W* positively regulates 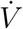 rather than the value of *V* (Fig. 1A). However, the relationship between the change in *W* and 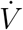 also does not reflect the positive regulation of *W* on *V*. For example, in the yellow region (Fig. 1B), 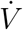 decreases despite increasing *W*, which happens because the self-regulation of *V* on 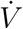 masks the effect of *W* on 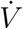. Thus, to infer the effect of *W* on 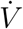 independently of *V*, we investigate the relationship between *W* and 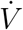 at the pair of time points *t* and the *reflection time, t*_*V*_, where *V* (*t*) = *V* (*t*_*V*_) (Fig. 1B). As *V* (*t*) = *V* (*t*_*V*_), the difference in 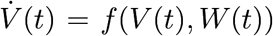 and 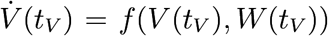 is solely determined by the difference between *W* (*t*) and *W* (*t*_*V*_). Thus, because *W* positively regulates *V* (Fig. 1A), if *W* (*t*) is greater (less) than *W* (*t*_*V*_), 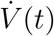 should be greater (less) than 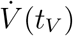. Similarly, to infer the type of self-regulation of *V*, we must remove the variation of 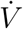 due to *W* that masks the effect of *V* on 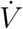. Thus, we investigate the relationship between *V* and 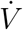 at the pair of time points *t* and the reflection time, *t*_*W*_, where *W* (*t*) = *W* (*t*_*W*_) (Fig. 1B). To quantify such relationships between *W* and 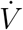 and *V* and 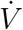, we develop the *regulation-detection functions*:

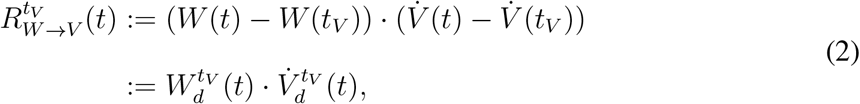

and

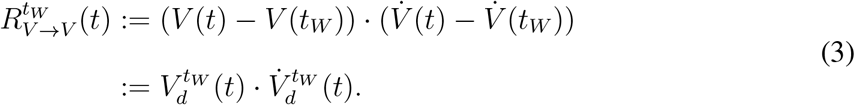

As *W* positively regulates *V*, the functions 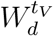 and 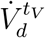 should have the same sign and thus, 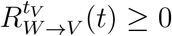 throughout the cycle (Fig. 1C, left). That is, if 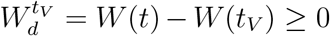, then 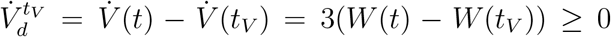 (Fig. 1A). On the other hand, due to the mixture of positive and negative self-regulation of *V*, the relationship between the signs of 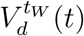 and 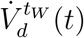, and thus the sign of 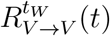, varies throughout the cycle (Fig. 1C, right). As the profiles of the sign of the regulation-detection functions (Eqns. (2) and (3)) reflect the regulation type, we develop a *regulation-detection score* that quantifies the variation in the sign of the regulation-detection functions. For instance, the regulation-detection score for the regulation of *W* on *V* is defined as

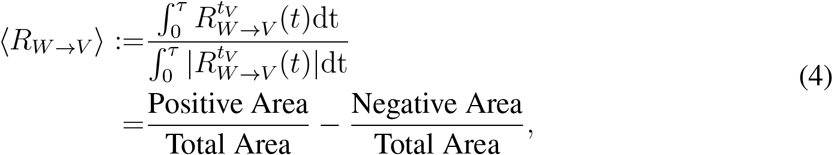

where *τ* is the period (e.g., *τ* = 1 in Fig. 1C, left). The regulation-detection score ⟨*R*_*W*→*V*_⟩ = 1 because *W* positively regulates *V*, and thus 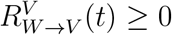 (i.e., the negative area is zero) (Fig.1C, left). On the other hand, because *V* both positively and negatively regulates itself, 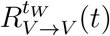 takes both positive and negative values, so ⟨*R*_*V*→*V*_ ⟩ = 0.6 − 0.4 = 0.2 (Fig. 1C, right).

Similarly, we can obtain information about the regulation of *V* on *W* and the self regulation of *W* with the regulation-detection functions 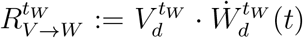 (Fig. 1D, left) and 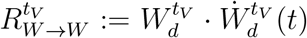 (Fig. 1D, right). Because *V* negatively regulates *W*, 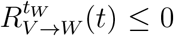. Also, because the self-regulation of *W* is purely negative, 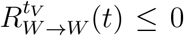. Thus, ⟨*R*_*V*→*W*_ ⟩ = −1, and *R*_*W*→*W*_ ⟩ = −1 (Fig. 1D). Taken together, in general, if *X* positively (negatively) regulates *Y*, then ⟨*R*_*X*→*Y*_ ⟩ = 1 (⟨*R*_*X*→*Y*_ ⟩ = −1) (see Theorem 1 in Supplementary Information).

Next, we calculated the regulation-detection scores from experimentally measured oscillatory time-series data of two bacteria: *Paramecium* and *Didinium*, which we refer to as P and D (Fig. 1E), respectively (*48*). As P is a prey of the predator D (*48*), D is expected to negatively regulate P, and P is expected to positively regulate D. Reflecting this, ⟨*R*_*P*→*D*_⟩ = 1 and ⟨*R*_*D*→*P*_ ⟩ = −1 (Fig. 1E). Furthermore, reflecting the positive (i.e., birth) and negative (i.e., death) self-regulation of both P and D, ⟨*R*_*D*→*D*_⟩ = 0.51 − 0.49 = 0.02 and ⟨*R*_*P*→*P*_⟩ = 0.63 − 0.37 = 0.26 (Fig. 1E). The regulation-detection scores appear to accurately reflect types of regulation even for noisy and discrete time-series data.

### Network Inference method from oscillatory time series

If *X* positively (negatively) regulates *Y*, then the reflection score ⟨*R*_*X*→*Y*_ ⟩ = 1 (resp., −1). In other words, −1 < ⟨*R*_*X*→*Y*_ ⟩ < 1 indicates either a mixture of positive and negative regulation of *X* to *Y* or the absence of regulation. Thus, in the system where the interactions are not mixed (i.e., monotonic), such as gene regulation by a transcription factor and predator-prey relationships, −1 < ⟨*R*_*X*→*Y*_ ⟩ < 1 indicates the absence of regulation. This can be used to infer network regulations from time-series data, as positive or negative regulation is present in the network only when ⟨*R*_*X*→*Y*_ ⟩ = 1 or −1, respectively. Similarly, self-regulation, which is either positive or negative, is possible only when ⟨*R*_*Y*→*Y*_ ⟩ = 1 or −1. However, since the degradation of molecules or the death rate of species typically increases as its own concentration increases, self-regulation can be assumed to be negative (i.e., ⟨*R*_*Y*→*Y*_ ⟩ = −1). In this case, positive or negative regulation from *X* to *Y* is possible only when 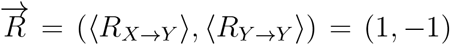 or (−1, −1), and thus, 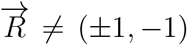 indicates the absence of regulation (Rule 1, Fig. 2A). Furthermore, we use 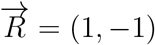 or (−1, −1) to infer positive or negative regulation (Rules 2 and 3, Fig. 2A). Note that, if positive or mixed self-regulation is possible, as in Fig. 1E, Rules 2 and 3 can be relaxed to ⟨*R*_*X*→*Y*_ ⟩ = 1 and ⟨*R*_*X*→*Y*_ ⟩ = −1, respectively.

**Figure 2:**
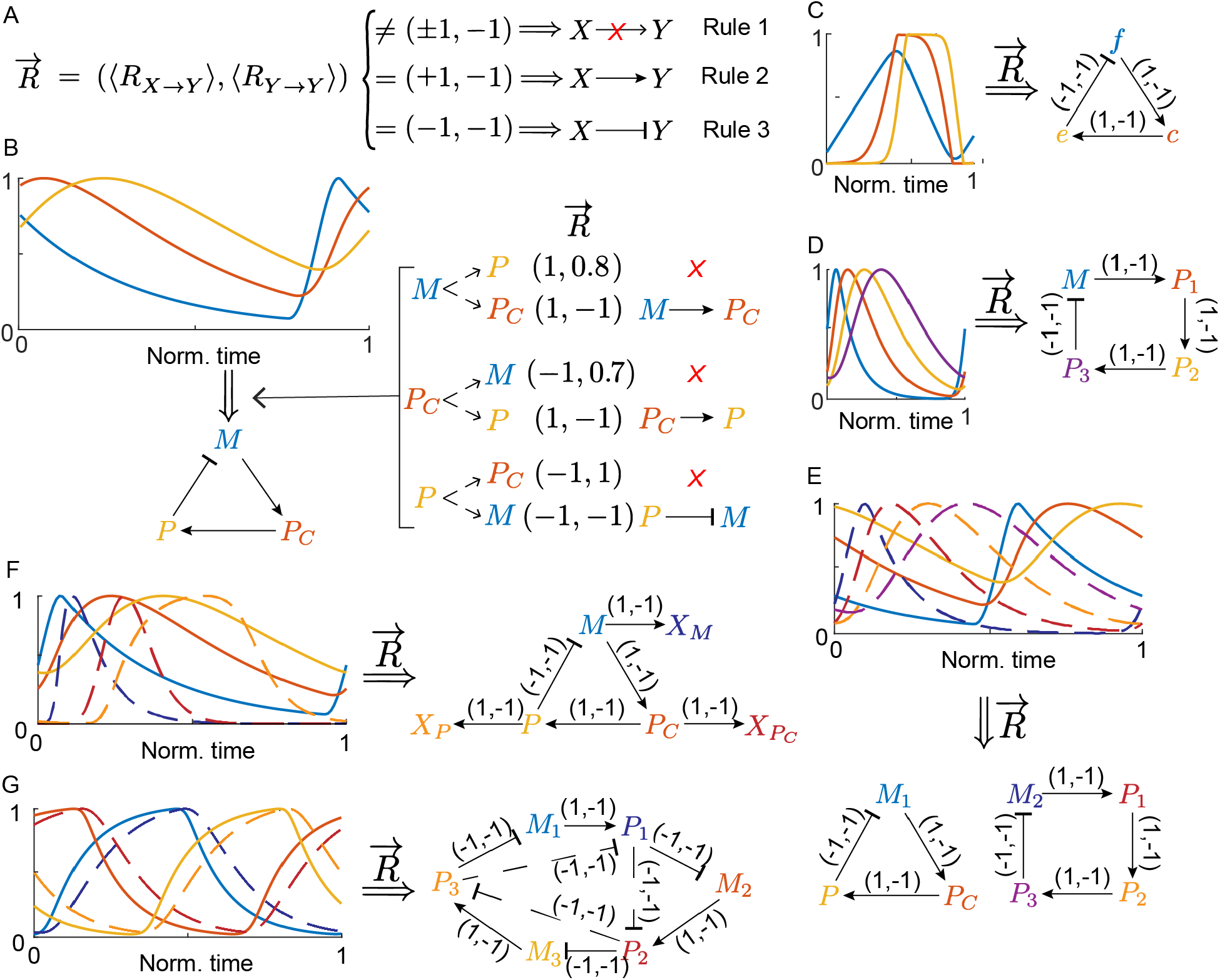
The inference method successfully infers various *in silico* network structures. (A) The three rules for network inference. 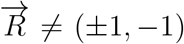 indicates the absence of regulation and 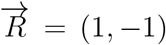 or (−1, −1) indicates positive or negative regulation. (B) The three rules successfully infer the network structure of the Kim-Forger model from simulated time-series data. According to Rule 1, the three regulations *M* → *P, P*_*C*_ → *M*, and *P* → *P*_*C*_ are inferred as absent. According to Rules 2 and 3, the two positive regulations (*M* → *P*_*C*_ and *P*_*C*_ → *P*) and the one negative regulation (*P* ⊣ *M*), which have 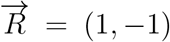 and 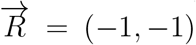, are inferred. (C-D) Our inference method also successfully infers the negative feedback loop of the *Frzilator* (C) and a 4-state Goodwin oscillator (D). (E-F) Our inference method also successfully infers correct regulations for more challenging cases beyond the single feedback loop structure, i.e., when time-series data are simulated with two independent models, the Kim-Forger model and Goodwin model (E) and an extended Kim-Forger model with output variables (F). (G) Our method also successfully infers regulations (solid arrows) of the repressilator from its three mRNA (solid lines) and three protein time-series data (dashed lines). However, our method also falsely predicts negative regulations among the proteins (dashed arrows) due to the similar time series between an mRNA and its protein (e.g., *M*_1_ and *P*_1_). See Tables S1-S5 for the complete list of regulation-detection scores for (C)-(G) and Section 3 in Supplementary Information for the equations and parameters used to simulate the data.

We illustrate how the three rules (Fig. 2A) can infer a network using as an example the Kim-Forger model (Fig. 2B), a simple model describing the transcriptional negative feedback loop of the mammalian circadian clock (*49, 50*). In the model, the mRNA (*M*) is translated into the cytosolic protein (*P*_*C*_). Then, *P*_*C*_ is transported to the nucleus and there *P* inhibits the transcription of *M* (*49,50*). To infer the network structure (Fig. 2B, bottom), we first compute 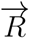 for each possible interaction and self-regulation pair (six in total) from the time series (Fig. 2B, top). Then, using Rule 1, three regulations are inferred as absent (Fig. 2B). Furthermore, Rules 2 and 3 successfully identify the two positive regulations (*M* → *P*_*C*_ and *P*_*C*_ → *P*) and the one negative regulation (*P ⊣ M*), which have 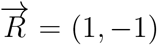 and 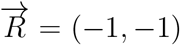, respectively. This successfully infers the negative feedback loop structure (Fig. 2B). Using the same procedure, our method also successfully infers the *Frzilator* negative feedback loop, which models the signaling circuit of *Myxococcus xanthus* (*51*) (Fig. 2C and Table S1) and a 4-state Goodwin oscillator (*52*) (Fig. 2D and Table S2).

In fact, for the single negative feedback loop models, the order of peaks and nadirs of the time series matches with the order of regulation in the feedback loop (Fig. 2B-D). For instance, the peak of *M* is followed by the peaks of *P*_*C*_ and then *P* (Fig. 2B). This property has been used in previous algorithms to infer single negative feedback loop structures (*36–38*). Next, we test whether our method can be applied to a more challenging case when data are merged from two independent models, specifically the Kim-Forger (Fig. 2E; solid lines) and Goodwin (Fig. 2E; dashed lines) models. After merging the time-series data, the order of peaks and nadirs cannot be used to infer the network anymore. That is, if only the order of peaks is used for this example, a single negative feedback loop with seven components is inferred. However, as our method uses the whole data set rather than just the peaks and nadirs, it successfully infers the two independent underlying networks (Fig. 2E, right and Table S3). Moreover, our inference method also successfully infers a cyclic network with output variables, which also does not adhere to the single feedback loop structure (Fig. 2F and Table S4).

While our method successfully infers various networks, Rules 2 and 3 can make false-positive inferences as 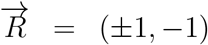 is a necessary condition for positive or negative regulation, and thus 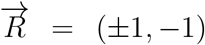 can occur even in the absence of regulation. We illustrate this using a simulated repressilator data set (Fig. 2G, left). The repressilator is a single feedback loop of genetic inhibition that consists of three mRNA (Fig. 2G, left; solid lines) and three proteins (Fig. 2G, left; dashed lines) (*53, 54*). The mRNA (*M*_*i*_) are translated into the respective proteins (*P*_*i*_), which then repress the transcription of the next gene (e.g., *P*_1_ represses *M*_2_). While our method recovers the correct interactions (Fig. 2G, right; solid arrows), it also incorrectly predicts negative regulation among the proteins (Fig. 2G, right; dashed arrows). These false-positive predictions are due to the similar shape and phase of the time-series data. For instance, the shape and phase of *P*_1_ (solid blue line) is extremely close to the shape and phase of its mRNA, *M*_1_ (dashed blue line), e.g., their phase difference is only 2.4% of the total period. Due to their similarity, our method cannot distinguish *M*_1_ and *P*_1_ and thus predicts that *P*_3_ negatively regulates not only *M*_1_ but also *P*_1_. For the same reason, our method falsely predicts that *P*_1_ negatively regulates *P*_2_, and *P*_2_ negatively regulates *P*_3_ (Fig. 2G and Table S5). Taken together, we caution that, in the presence of nearly identical time series in a network, our method may infer false-positive regulations, which seems unavoidable for any inference methods using time-series data.

### Robustness of the inference method to interpolation error and noise

The calculation of 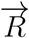, which is the key to the inference method, requires continuous time-series data. Typically, however, experimentally measured time-series data are sampled discretely. For instance, mRNA levels of circadian genes frequently are measured via PCR every three hours (*55*). For discrete data, our method uses interpolation to generate continuous data (see Methods). Accordingly, we test how sensitive our method is to interpolation error, specifically when linear interpolation is used, by using the five *in silico* data sets in Figs. 2B-F. That is, by decreasing the points measured per period from 10^2^ to 10^1^ (i.e., increasing the interpolation error), we quantify the accuracy of our network inference method with the score 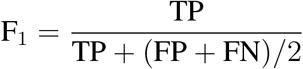 (TP-the number of true positives, FP-false positives, and FN-false negatives). As F_1_ is the harmonic mean of precision and recall, F_1_ = 1 and F_1_ = 0 indicate perfect recovery of the network and absence of correct inference, respectively. To account for interpolation error, we accept interactions based on three thresholds for ⟨*R*⟩ values: 0.99, 0.95, and 0.90. For example, a threshold of 0.99 means that we relax the condition 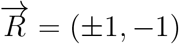 up to ±0.99, i.e, we accept any interaction that satisfies both |⟨*R*_*X*→*Y*_ ⟩| > 0.99 and ⟨*R*_*Y*→*Y*_ ⟩ < −0.99. We repeat this process 100 times, each time beginning the sample collection at a randomly selected time in the period (see Methods for details). Then, we investigate how the mean of the distribution of F_1_ scores changes as the sampling rate decreases (Fig. 3A). For single negative feedback loops (i.e., Frzilator, Goodwin, Kim-Forger), our method accurately recovers the network even when the number of data points measured per period is relatively low, e.g., ten per cycle. For the more complicated models (i.e., the merged Goodwin and Kim-Forger and the Kim-Forger with outputs models), slightly more data points are required for inference at high accuracy. Furthermore, our method shows similar robustness across the three thresholds, especially when the points sampled are toward the lower end.

**Figure 3:**
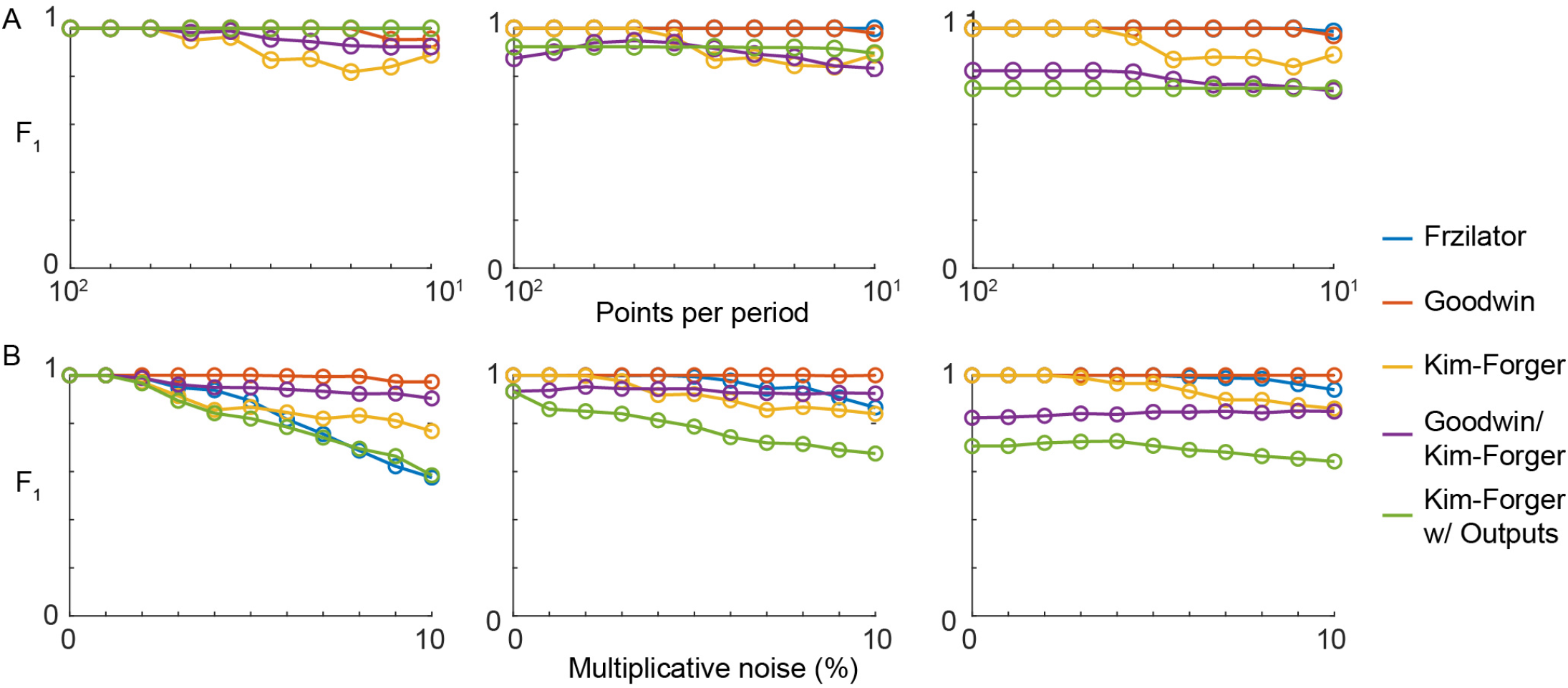
Our network inference method is robust to interpolation error and to noise. (A-B) The accuracy of our inference method when the number of points measured (A) and the level of noise (B) vary. Here, the points measured per period decreases from 10^2^ to 10^1^ (A) and the multiplicative noise increases from 0 to 10%, which is sampled from N(0, 0.1^2^) (B). The mean of the F_1_ score for 100 different time series, which are generated with randomly chosen phases and noise levels (B), is plotted (see Methods for details). F_1_ = 1 and 0 indicate perfect recovery of the network and the absence of correct inference, respectively. Different thresholds for ⟨*R*⟩, 0.99 (left), 0.95 (middle), and 0.90 (right), are used.

Next, because experimental data includes noise, we test the sensitivity of our network inference procedure to noise (Fig. 3B) (see Methods for details). As we increase the level of the multiplicative noise added to the data set from 0 (no noise) to 10% multiplicative noise (sampled from N(0, 0.1^2^)), the F_1_ scores decrease. In particular, the decrease occurs more dramatically when the threshold is 0.99, indicating that the high threshold leads to higher sensitivity to noise in the data. Moreover, this decrease in F_1_ scores with the threshold of 0.99 is a result of an increase in false negatives (i.e., the exclusion of true interactions due to noise). Thus, we use a threshold of 0.9 when applying our inference method to experimental data (see below) as it leads to the most accurate results in the presence of noise (Fig. 3B). However, users have the option to adjust the threshold depending on the sampling rate and noise level of the data when using our computational package, ION (Fig. 4A) see Supplementary Information and Figs. S1 and S2 for a step-by-step manual).

**Figure 4:**
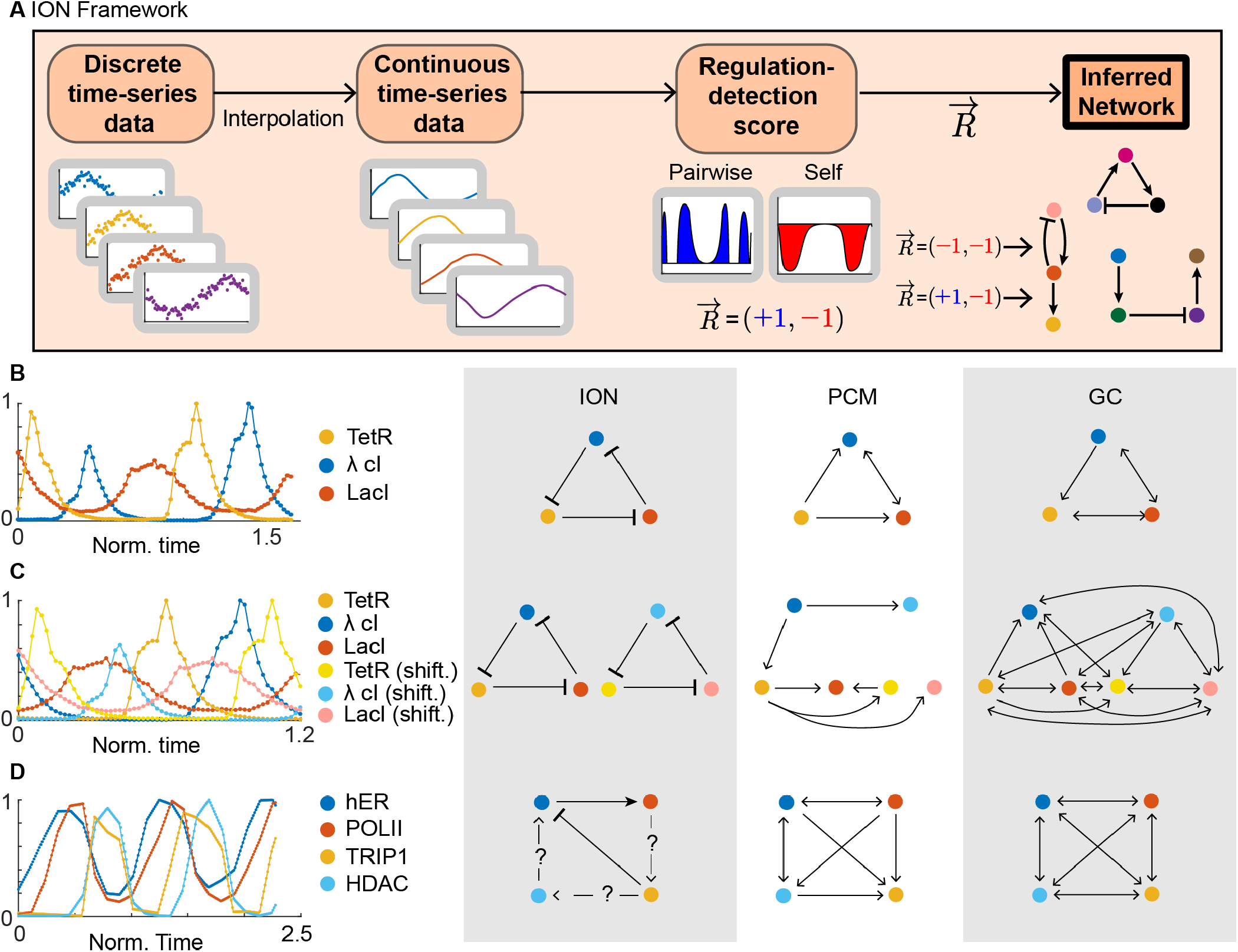
The computational package ION successfully infers networks of repressilator and pS2 promoter cofactors. (A) The ION package interpolates and smooths noisy and discrete time-series data and then computes the pairwise and self regulation-detection functions and scores 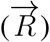 for every pair of components. If 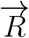 satisfies Rules 2 or 3 (Fig. 2A), which can be relaxed depending on the threshold specified by the user, positive or negative regulation is assigned. See Supplemental Information for a comprehensive manual. (B) Using experimentally measured oscillatory time series from (*54*), ION successfully infers a three-gene repressilator network structure. On the other hand, two popular model-free inference methods, PCM and GC, infer several false-positive regulations (e.g., *λ* cI regulates LacI). (C) ION also successfully infers two independent cycle when the experimental repressilator data set from (A) is duplicated and the phase is shifted by about half of the period. However, PCM and GC fail in infring independent cycles due to false-positive predictions. (D) ION infers two direct regulations and predict three hidden regulations among cofactors at the estrogen-sensitive pS2 promoter after estradiol treatment (*57*).

### Successful inferences from experimentally measured time series

As our inference method is quite robust to discrete data sampling and noise, we expect that our inference method can accurately infer network structures from experimentally measured time series as well. Indeed, when applied to experimentally measured abundances of the three repressilator proteins (*54*), our method successfully infers a three-gene repressilator network structure (Fig. 4D and Table S6). Note that our method recovers the repressilator network despite the absence of mRNA data because the shape and phase of the mRNA and protein profiles are expected to be similar, as in Fig. 2G, due to the short translation time in *E. coli* compared to the period (*56*). Moreover, we compare the results of our method with those of two popular model-free inference methods, Partial Cross Mapping (PCM) (*20*) and Granger Causality (GC) (*4*) (Fig. 4D). As these methods can only infer the presence of regulation, not its type (i.e., positive and negative), unlike our method, the arrows represent inferred regulations, which could be either positive or negative. The PCM method recovers two correct regulations, *P*_2_ → *P*_1_ and *P*_3_ → *P*_2_, but fails to recover the regulation *P*_1_ → *P*_3_ and makes two false-positive predictions, *P*_1_ → *P*_2_ and *P*_3_ → *P*_1_. While the GC method infers all existing regulations, it makes two additional false-positive predictions, *P*_1_ → *P*_2_ and *P*_2_ → *P*_3_. Even for this simple three-node network, the popular model-free inference methods make false-positive predictions because the network components oscillate at the same period.

We compare the performance of our method with these model free-inference methods for a more challenging case when we combine two copies of the data set in Fig. 4D, one at the original phase and one with shifted phase (Fig. 4C and Table S7). From the combined time-series data, our method successfully infers two repressilator networks, whereas the PCM method infers two of the six correct regulations while also inferring four incorrect regulations (Fig. 4C). The GC method infers more correct regulations (five of the six); however, again, it suffers from several spurious regulations (15 false-positive interactions, Fig. 4C). Note that, even though we are using the same repressilator data set, there are inconsistencies in the PCM and GC results compared with those from Fig. 4D. These inconsistencies are a consequence of the shortened length of data used in Fig. 4C compared with that in Fig. 4D. This indicates that, in addition to the risk of false-positive inference, the PCM and GC methods are sensitive to the amount of data, unlike ours.

For time series measuring the amount of cofactors present at the estrogen-sensitive pS2 promoter after treatment with estradiol (data from (*57, 58*)), PCM and GC infer an almost fully connected network and a fully connected network, respectively (Fig. 4D). On the other hand, our method only infers two regulations, both supported by the current biological understanding of the system. That is, human ER*α* (hER) binds to the pS2 promoter after treatment with estradiol to recruit RNA Polymerase II to the promoter, supporting the inferred positive regulation of POLII by hER. Furthermore, TRIP1 acts as a surrogate measure for the 20S proteasome (APIS), which promotes proteasome-mediated degradation of hER (*57*), supporting the inferred negative regulation of hER by TRIP1. However, the inferred network (Fig. 4D, Table S8) does not contain a negative feedback loop, which is required to generate sustained oscillations (*59*). Thus, there may be intermediate steps between POLII and TRIP1, TRIP1 and HDAC, and also HDAC and hER that form the negative feedback loop. Altogether, this illustrates that our method can identify direct regulations while highlighting connections that require further experimental investigation.

## Discussion

We developed a model-based method that infers the network structure underlying biological oscillators. The method identifies positive or negative regulation by testing whether given oscillatory time-series data are reproducible with a general mechanistic ODE model (Eqn. (1)). In this way, our method successfully and efficiently inferred several network architectures such as single cycles (e.g., repressilator), two independent cycles, and a cycle structure with outputs. Furthermore, we provide a user-friendly computational package, ION, that applies to discrete and noisy data to infer networks of biological components that oscillate from the molecular to the population level. Our method can uncover unknown functional relationships and mechanisms that drive oscillatory behavior in biological systems when it is incorporated with evolving experimental time-series measurement methods.

Our method merges the advantages of model-based and model-free methods while mitigating the drawbacks of each. In particular, our model-based inference method does not suffer from the serious risk of false-positive prediction for biological oscillators or sensitivity to the amount of data unlike the previous model-free inference methods such as GC and PCM (Fig. 4). However, as our inference method is model-based, it runs the risk that the imposed ODE model and functional relationships create false representations of the true interactions (*21*). Our method minimizes this risk by using the most general form of an ODE (Eqn. (1)) to model the change in a component that is acted upon by another component and itself. In this way, we resolve the limitations of previous model-based methods that restricted the class of models, such as separable synthesis and degradation functions (*39, 41, 45*), specific types of functions (e.g., power or Hill functions) (*31, 39*), and a single negative feedback loop structure (*36–38*). Thus, we were able to uncover several varying network structures. While we considered the most general form of an ODE (Eqn. (1)) that describes the interactions between two components, an interesting future direction would be to extend our work to models that describe the interactions among multiple oscillatory components, e.g, 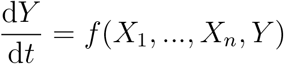.

## Methods

### ION (Inferring Oscillatory Networks) computational package

We provide user-friendly MATLAB code (a Github repository link will be provided upon acceptance of the manuscript). The ION package can be used to infer the network structure of oscillators, which are described by Eqn. (1), across all levels of biology. Here, we briefly describe the key steps of the ION package (see Supplementary Information for a comprehensive manual).

#### Reflection times

For each time point *t*_*i*_ of the given time series *X*(*t*) = (*X*(*t*_1_), *X*(*t*_2_), …, *X*(*t*_*n*_)), first, the reflection time *t*_*iX*_ needs to be calculated (Fig. 1B). That is, we seek the time point *t*_*iX*_ such that *X*(*t*_*i*_) = *X*(*t*_*iX*_) and the signs of the slopes at *X*(*t*_*i*_) and *X*(*t*_*iX*_) are opposite (i.e., rising and falling phase). For this, the discrete *X*(*t*) is interpolated to a continuous time series *F*_*X*_(*t*) with either the ‘linear’ or ‘fourier’ interpolation method, chosen by the user. Then, *t*_*iX*_ is estimated by identifying the closest time point to *t*_*i*_ among time points *t* satisfying the following equation:

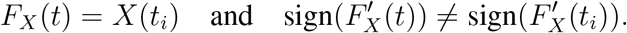

#### Regulation-detection function and score

Using the estimated *t*_*iX*_, we compute the regulation-detection function, e.g., 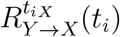, for each time point *t*_*i*_ as follows:

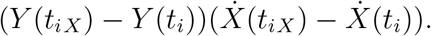

If the linear method is chosen, *Y* (*t*_*iX*_) is linearly interpolated based on the data (*Y* (*t*_1_), …, *Y* (*t*_*n*_)), and 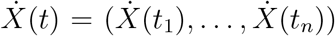 is estimated using a moving slope filter method. Specifically, after fitting a low-order polynomial regression model to *X*(*t*) = (*X*(*t*_1_), *X*(*t*_2_), …, *X*(*t*_*n*_)) in a sliding window (*60*), the derivative of the polynomial fit is used to estimate 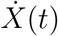 and then 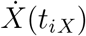 is linearly interpolated based on the estimated 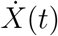. The model order and the length of the sliding window parameters can be adjusted (see Supplementary Information). On the other hand, if the fourier method is chosen, both 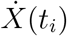 and 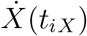 are estimated as 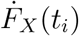 and 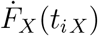, respectively, and similarly, *Y* (*t*_*i*_) and *Y* (*t*_*iX*_) are estimated as *F*_*Y*_ (*t*_*i*_) and *F*_*Y*_ (*t*_*iX*_), respectively, where *F*_*Y*_ (*t*) is the Fourier series fit to the data *Y* (*t*). Finally, in both cases, the regulation-detection score Eqn. (4) is estimated using the MATLAB function `trapz`.

#### Time-series data

We simulate *in silico* data using the MATLAB function `ode23tb` (Fig. 2). See Supplementary Information for the model equations and parameters. The experimental data sets of the repressilator (Fig. 4D) were obtained from (*54*). Next, to generate the duplicated experimental repressilator data set (Fig. 4C), we mixed two copies of the repressilator data set from Fig. 4D. We kept one copy at the original phase and, for the second copy, we shifted the phase by 115 minutes (almost half of the period) (Fig. 4C). Then, we removed data on the left and the right where there was only coverage of one of the two data sets. We obtained the estradiol data set from (*57, 58*) and the Paramecium/Didinium data from (*10*).

#### Discrete and noisy data

To generate discretely sampled data (Fig. 3A), we select a random point in the first period to begin data extraction, and then we uniformly sample two periods worth of data at a sampling rate of 100 points per period. We repeat this process 100 times–every time randomly initializing the starting point in the first period–to generate 100 distinct data sets for every model. Then, we run our algorithm and compute F_1_ scores for each of the 100 data sets. Next, from each of the 100 generated data sets, we take every other data point to reduce the number of data points (e.g., 50, 33, 25, …, 10 per period).

For the multiplicative noise analysis (Fig. 3B), we begin with two periods worth of data sampled at 100 points per period. Then, we add multiplicative noise sampled randomly from a normal distribution with mean 0 and standard deviation given by the percentage. For example, at 10% multiplicative noise, we add the noise *X*(*t*_*i*_) *· ϵ* to *X*(*t*_*i*_), where *ϵ* is sampled randomly from N(0, 0.1^2^).

#### PCM and GC

We ran the PCM method with an embedding dimension of 3, *τ* = 1, and a max delay of 3, and used a threshold of 0.5684 as recommended in (*20*) by using the code provided in (*20*). We ran the GC using the code provided in (*61*) and specified a max delay of 3 as we did with the PCM method and a significance level of 95%. We rejected the null hypothesis that *Y* does not Granger cause *X*, and thereby inferred direct regulations if the value of the F-statistic was greater than the critical value from the F-distribution (*4*).

## Acknowledgments

We thank Anne Shiu and Seokjoo Chae for valuable comments. This work was supported by a National Institutes of Health Training Grant (T32 HL007622) and Samsung Science and Technology Foundation under Project Number SSTF-BA1902-01 (to J.K.K.).

## Author Contributions

JT, DF, and JKK designed the research. JT and JKK conceptualized the theoretical approach, performed the research, and developed the computational platform. JT, DF, and JKK wrote the manuscript.

## Supplemental Information

### 1 Theoretical Foundations

Here, we describe the theoretical foundations of our main result. In particular, we formally define the reflection time *t*_*Y*_ (Fig. 1B) and prove that, if *X* positively regulates *Y*, then the regulation-detection score ⟨*R*_*X*→*Y*_⟩ = 1. Similar arguments can be used for negative regulations.

#### Definition 1.

*If Y* (*t*) *is a smooth time series with one maximum (t*_*M*_ *) and one minimum (t*_*m*_*) over a period, then for t* ≠ *t*_*M*_, *t*_*m*_, *we define the reflection time t*_*Y*_ *such that Y* (*t*_*Y*_) = *Y* (*t*) *and t*_*Y*_ ≠ *t. If t* = *t*_*M*_, *t*_*m*_, *then t*_*Y*_ := *t*.

#### Theorem 1.

*Let X*(*t*) *and Y* (*t*) *be smooth time series with period τ that have one maximum and one minimum over a period τ and solve the ODE*

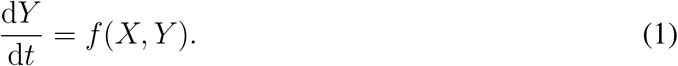

*If f is strictly monotonically increasing in X (i*.*e*., *X positively regulates Y), then* 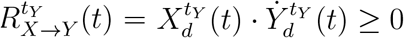 *for all t, where*

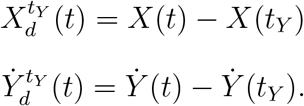

*Moreover,* ⟨*R*_*X*→*Y*_⟩ = 1, *where*

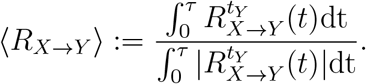

*Proof*. First,

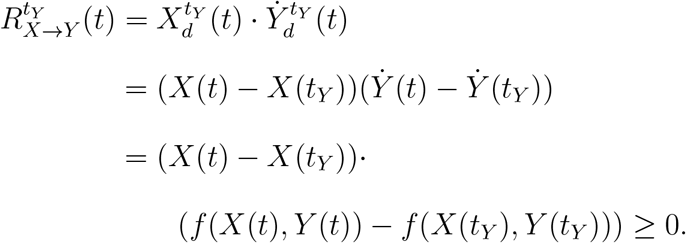

The last inequality holds because, by assumption, *f* (*X, Y*) is a monotonically increasing function of *X* when *Y* has the same value (i.e., *Y* (*t*) = *Y* (*t*_*Y*_)). Furthermore, 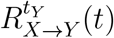 cannot be 0 for all *t*. For the sake of contradiction, assume that 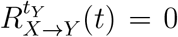 for all *t*. If 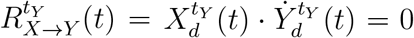, then 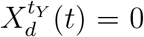 or 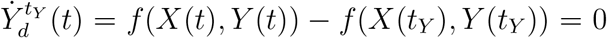. If *f* (*X*(*t*), *Y* (*t*)) − *f* (*X*(*t*_*Y*_), *Y* (*t*_*Y*_)) = 0, then *X*(*t*) − *X*(*t*_*Y*_) = 0 as well because *f* is strictly monotonically increasing in *X*. Therefore, if 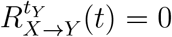 for all *t*, then

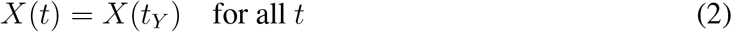

Next, from (1) and (2),

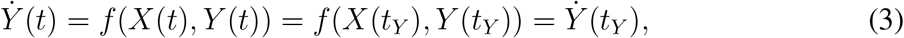

for all *t*. This implies that 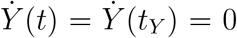 given the nature of *t*_*Y*_ (e.g, if *Y* is increasing at *t*, then *Y* is decreasing at *t*_*Y*_). Therefore, 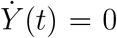 for all *t*, i.e., *Y* is constant. However, this contradicts the assumption that *Y* is a smooth oscillating time series. Since 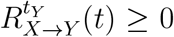 and is a nonzero function, ⟨*R*_*X*→*Y*_ = 1⟩.

### 2 Manual for the ION computational package

As described in the main text, we developed a computational package, ION (Inferring Oscillatory Networks), to infer networks of biological oscillators (Fig. 4A). We have published the ION package in MATLAB at the link: (a Github repository will be provided upon acceptance of the manuscript). ION can be used as follows.

1. *Generate the data*.*csv file*. The first column of the data.csv file should be the time points at which the measurements were taken. The subsequent columns should be the data for each variable at the respective time points. If the data are stored in an excel spreadsheet, order the file as time points in column 1, data for variable 1 in column 2, data for variable 2 in column 3, etc., and then save it as a .csv file in the same directory as the code. Be sure that there are no variable names at the top of the data.csv file (see Input 1 in Fig. S1 for the structure of the data.csv file).
2. *Update the* `regulation_detection_inputs.m` *file*. Users need to specify the threshold used to infer regulation and the data interpolation method, in particular, either ‘linear’ or ‘fourier’. The linear method will linearly interpolate the discrete data given in the data.csv file to create a continuous data set. The fourier method will fit a Fourier series to the discrete data to create a continuous data set.
  a. *Update the threshold variable* (Input 2 in Fig. S1). The variable *threshold* determines the threshold for the 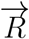 values accepted as interactions. For example, a threshold of 0.99 means that we relax the condition 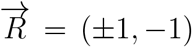 (Fig. 2A) up to *±*0.99, i.e, we accept any interaction that satisfies both |⟨*R*_*X*→*Y*_ ⟩| *>* 0.99 and ⟨*R*_*Y* →*Y*_ ⟩*< −*0.99. Based on our analysis in Fig. 3, we recommend a threshold of 0.9, which we use for inferring interactions given experimental data in Fig. 4.
  b. *If linear method is chosen, update the variables ‘supportlength’ and ‘modelorder’* (Input 2 in Fig. S1). They need to be chosen so that the function `movingslope.m` accurately estimates the derivatives at each time point of the data set (*1*). The variable ‘supportlength’ determines the number of points used for the moving window average estimation of the derivative, so it should be at least 2 and no more than the number of data points. If ‘supportlength’ is odd, then the derivative is estimated using a central sliding window. If ‘supportlength’ is even, then the moving window will be moved backward by one element. For example, a ‘supportlength’ of 2 is equal to a difference quotient, computed for instance using the MATLAB function `diff`. Next, the variable ‘modelorder’ defines the order of the windowed model used to estimate the slope. If ‘model order’ is 1 or less than the ‘supportlength’-1, then the model is linear or a regression, respectively. If ‘modelorder’ is equal to ‘supportlength’-1, then the method will be a sliding Lagrange interpolant.
  c. *If fourier method is chosen, update the ‘num_fourier’ variable* (Input 2 in Fig. S2). The variable ‘num_fourier’ determines the number of Fourier series coefficients and must be at least one and at most 8. For example, if ‘num_fourier’ = 1, then the algorithm fits the parameters *a*_0_, *a*_1_, *b*_1_, and *ω* to the data for each variable based on the Fourier series of the form *F* (*t*) = *a*_0_ + *a*_1_ cos(2*π* · *ω* · *t*) + *b*_1_ sin(2*π · ω · t*). The algorithm fits the Fourier series with the specified order using a nonlinear least-squares method, specifically using the Levenberg-Marquardt method. To manually adjust the Fourier fitting method and options (e.g., the method, function tolerance, etc.), users need to update the function `createFit.m`.
3. *If linear method is chosen, create extrema value files for each variable and save them in the same directory as the scripts* (Input 3 in Fig. S1). If the fourier method is chosen, the interpolated time series has one maximum and minimum per cycle. However, if the linear method is chosen, local extrema can occur due to noise. To distinguish such local extrema from the global extrema per cycle, which is required to compute the reflection time (see Fig. 1B), users need to input extrema value files for each variable in the data set. The files should be saved as e*k*.csv where *k* corresponds to the variable. For example, the extrema value file for variable 1 (the first data column) should be saved as e1.csv. The extrema value files should have time points in the first column and either 1 or −1 in the second column for a global maximum and a global minimum in that period at the time point, respectively. Users can generate these files by using the MATLAB function `findpeaks`. The function `findpeaks` returns all local minima and maxima, which users can use to decide the global minimum and maximum in each period.
4. *Run the* `main.m` *function*. The `main.m` function will read the inputs from the `regulation_detection_inputs.m` file and run the algorithm with the user-specified method. If the linear (fourier) method was chosen, the `main.m` function calls the `main_linear.m` (`main_fourier.m`) script to run the algorithm. Users will see several figures being generated as well as several output files generated in a new “Outputs” directory, which will be created within the same directory that the script is run:
  a. If the linear method is chosen, a derivative.fig file that plots the estimated derivative values for all variables (Output 1 in Fig. S1).
  b. If the fourier method is chosen, files labeled Var*k*.fig, where *k* corresponds to the variable number, showing the fourier fits for each variable (Output 1 in Fig. S2).
  c. Figures plotting the regulation-detection functions for each possible interaction (Output 2 in Fig. S1 and S2). For example, the file Reg_detect_i_onto_i_fix_j.fig plots the regulation-detection function 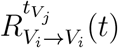 (see Fig. 1C), where *V*_*i*_ is variable *i*. Similarly, the file Reg_detect_i_onto_j.fig plots the regulation-detection function 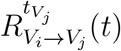 (see Fig. 1C).
  d. A .mat file with the regulation-detection scores for each possible self-regulation (‘R_self’ variable) and cross-regulation (‘R_cross’ variable) (Output 3 in Fig. S1 and S2).
  e. A figure plotting the inferred network structure, inferred_network_graph.fig (Output 4 in Fig. S1 and S2). If R_self(*i, j*) *<* −threshold and —R_cross(*i, j*)| > threshold, then the algorithm infers the interaction from Variable *j* to Variable *i*. Moreover, if R_cross(*i, j*) is positive (negative), then it is a positive (negative) regulation.
  f. A Final_results.pdf document that reports each output listed above for reference.

**Figure S1:**
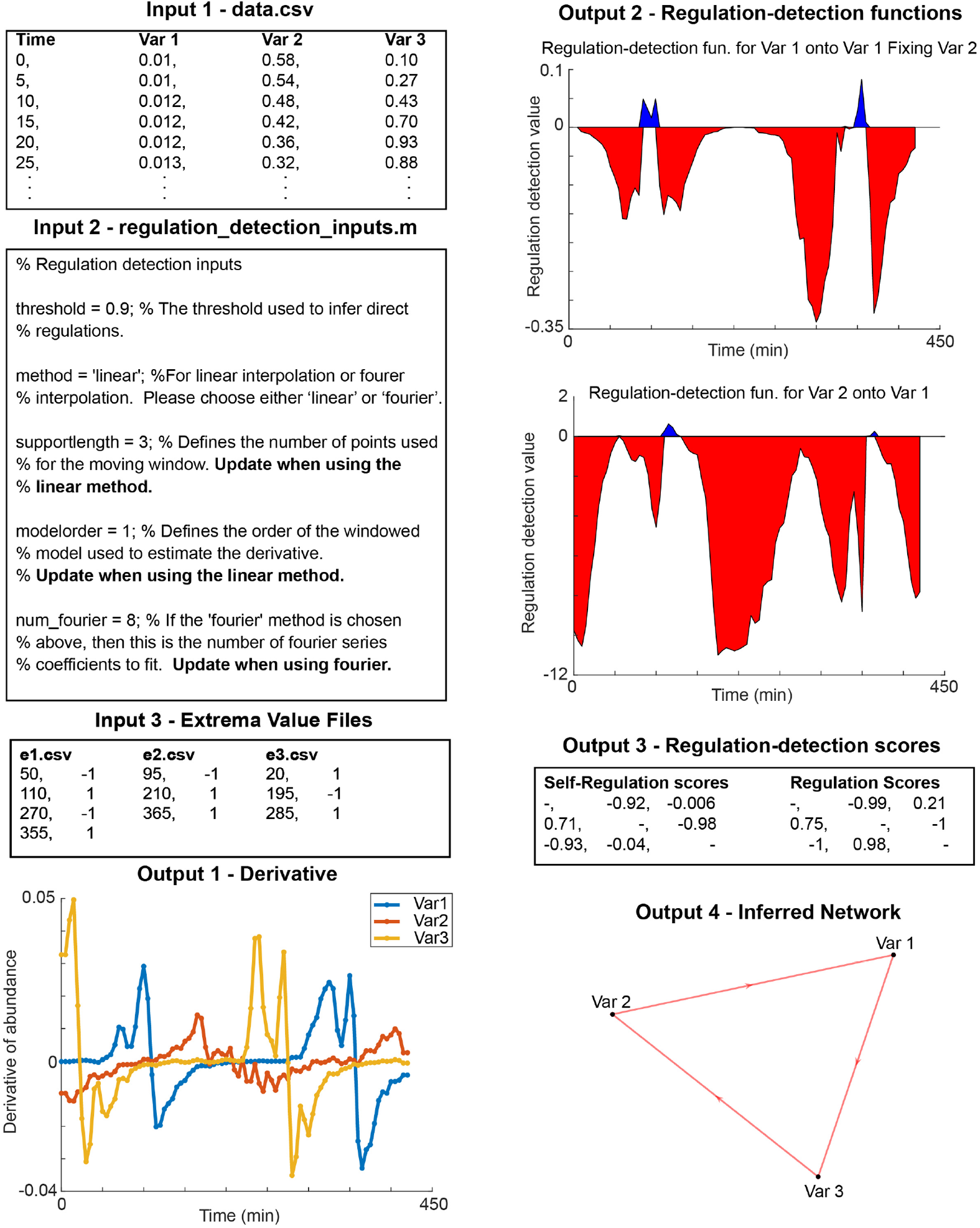
Sample input and output files for the ION package if the linear method is chosen based on the experimental repressilator example (Fig. 4B). The data.csv (Input 1) file contains the time points in the first column and the data for each variable in subsequent columns, separated by commas. The regulation_detection_inputs.m file (Input 2) is already in the directory and should be updated according to the preferences of the user. If the user selects the ‘linear’ method, then the user must choose appropriate values for the ‘supportlength’ and ‘modelorder’ variables as well as generate extrema value files that list the time points at which the global maxima (1) and minima (−1) happen for each period (Input 3). The algorithm outputs the estimated derivatives for each variable (Output 1), plots of the regulation-detection functions (Output 2), the regulation-detection scores (Output 3), and the inferred network structure (Output 4).

### 3 Description of the *in silico* models

#### 3.1 Kim-Forger Model

The Kim-Forger model describes the core transcriptional-translational feedback loop in the molecular circadian clock (Fig. 2B) (*2*). It is similar to the Goodwin model in that a protein product represses the transcription of its *mRNA*. However, the repression mechanism is not a Hill-type repression mechanism as in the Goodwin oscillator, but rather a protein sequestration mechanism where tight binding of the activator and repressor sequesters the activator protein, which in turn represses its activity. The model is given by the following system of ODEs.

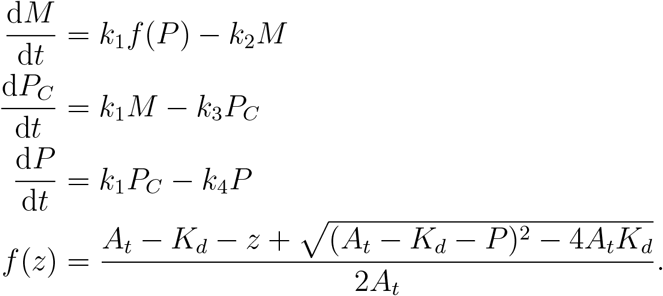

Here, the variable *M* is the repressor mRNA concentration; *P*_*C*_ is the cytosolic repressor protein concentration; *P* is the nuclear repressor protein concentration. The parameter *A*_*t*_ is the total activator concentration, and *K*_*d*_ is the dissociation constant between the activator and repressor (closer to 0 means tighter binding). As in (*3*), we simulate the Kim-Forger model with the parameters *k*_1_ = 1, *k*_2_ = 0.16, *k*_3_ = 0.29, *k*_4_ = 0.3, *A*_*t*_ = 0.6, and *K*_*d*_ = 10^−5^.

**Figure S2:**
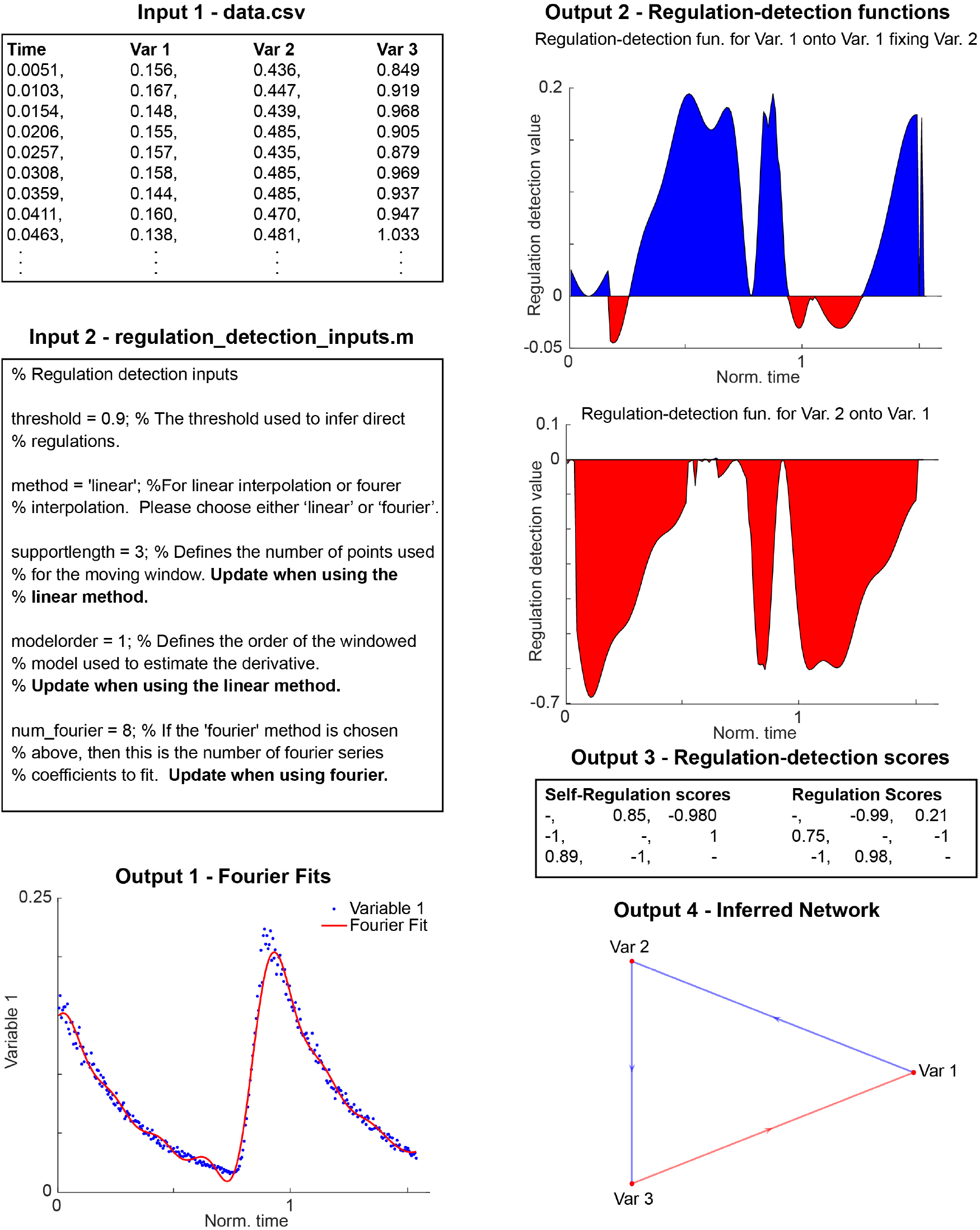
Sample input and output files for the ION package if the fourier method is chosen based on the Kim-Forger model with added noise (Fig. 2B). The data.csv (Input 1) file contains the time points in the first column and the data for each variable in subsequent columns, separated by commas. The regulation_detection_inputs.m file (Input 2) is already in the directory and should be updated according to the preferences of the user. If the user selects the ‘fourier’ method, then the algorithm fits a Fourier series to each variable with the number of coefficients specified by the variable ‘num_fourier’ variable (Input 2). The algorithm outputs the Fourier fits for each variable (Output 1), plots of the regulation-detection functions (Output 2), the regulation-detection scores (Output 3), and the inferred network structure (Output 4).

#### 3.2 Frzilator

The *frzilator* model consists of a negative feedback loop that models the oscillations in a cascade of covalent modifications in *Myxococcus xanthus* (*4*) (Fig. 2C). The mathematical model uses Michaelis-Menten dynamics to model the negative feedback and is given by the equations

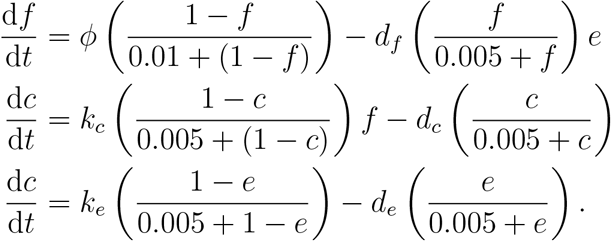

As in (*3*), we simulate the system with the nominal parameter values *ϕ* = 0.08, *k*_*c*_ = 4, *k*_*e*_ = 4, *d*_*f*_ = 1, *d*_*c*_ = 2, and *d*_*e*_ = 2.

#### 3.3 Goodwin Oscillator

The Goodwin model describes the action of a protein product repressing its *mRNA* (*5*) (Fig. 2D). The transcriptional regulation is described by a Hill function, where the Hill coefficient is an upper bound for the number of repressor proteins that bind to the promoter (*6*). A high Hill coefficient is required for the system to generate sustained oscillations. We simulated the Goodwin oscillator from the following system of ODEs.

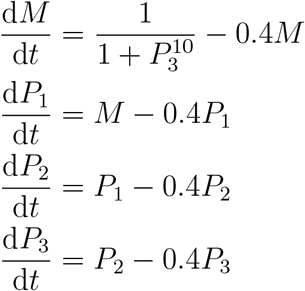

#### 3.4 Kim-Forger Model with Outputs

We augment the original Kim-Forger model (Section 3.1) so that each component promotes the production of one output, e.g., *M* positively regulates *X*_*M*_ (Fig. 2F). The system of ODEs is given below.

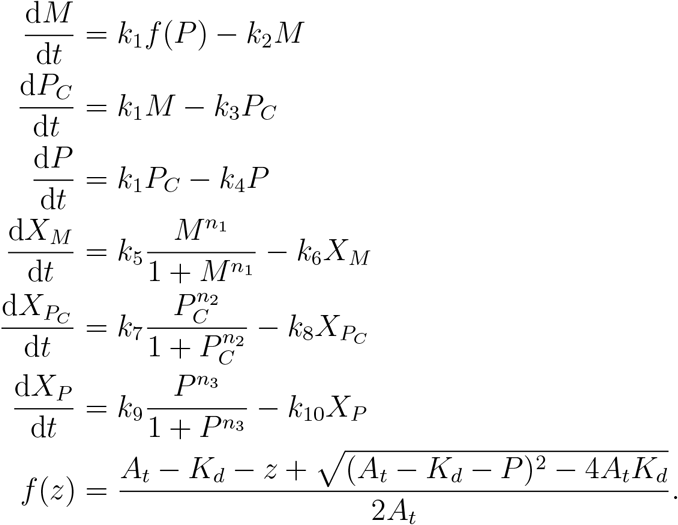

Here, the parameters *k*_1_ through *k*_4_ are the same as in Section 3.1. We simulate the model with the parameters *k*_5_ = 0.5, *k*_6_ = 0.59, *k*_7_ = 0.23, *k*_8_ = 0.98, *k*_9_ = 0.08 and *k*_10_ = 0.63 and Hill coefficients *n*_1_ = 10, *n*_2_ = 15, and *n*_3_ = 20.

#### 3.5 Repressilator

The repressilator is a synthetic feedback loop that consists of three genes and three proteins where the mRNAs translate to the respective proteins, which in turn repress the transcription of the next mRNA (Fig. 2G) (*7*). The model is given by the following ODEs.

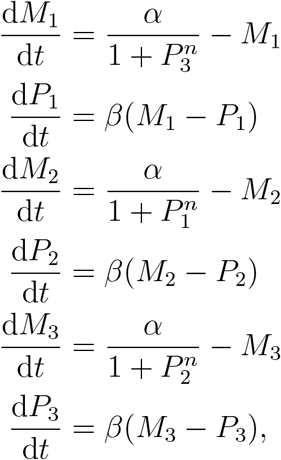

where *α* = 7, *β* = 3, and *n* = 10.

### 4 Complete results of *in silico* and experimental network inference (Figs. 2 and 4)

In Tables S1-S5, we report the complete results of our network inference procedure applied to the *in silico* examples in Fig. 2. In Tables S6-S8, we report the complete results of our network inference procedure applied to the experimental data sets (Fig. 4). The inferred interactions are the interactions that pass Rules 1-3 (Fig. 2A) with a threshold value of 0.90. Each table cell is (⟨*R*_*X*→*Y*_⟩, ⟨*R*_*Y* →*Y*_⟩) where *X* is the column variable and *Y* is the row variable.

**Table S1:** Calculated regulation-detection scores for the Frzilator oscillator example (Fig. 2C).

| | $f$ | $c$ | $e$ |
| --- | --- | --- | --- |
| $f$ | – | $(-1, 1)$ | $(-1, -1)$ |
| $c$ | $(1, -1)$ | – | $(-1, 0.99)$ |
| $e$ | $(1, -0.71)$ | $(1, -1)$ | – |

**Table S2:** Calculated regulation-detection scores for the Goodwin oscillator example (Fig. 2D).

| | $M$ | $P_1$ | $P_2$ | $P_3$ |
| --- | --- | --- | --- | --- |
| $M$ | – | (–1, 0.98) | (–1, 0.61) | (–1, –1) |
| $P_1$ | (1, –1) | – | (–1, 0.92) | (–1, 0.21) |
| $P_2$ | (1, –0.30) | (1, –1) | – | (–1, 0.64) |
| $P_3$ | (1, 0.06) | (1, –0.23) | (1, –1) | – |

**Table S3:**
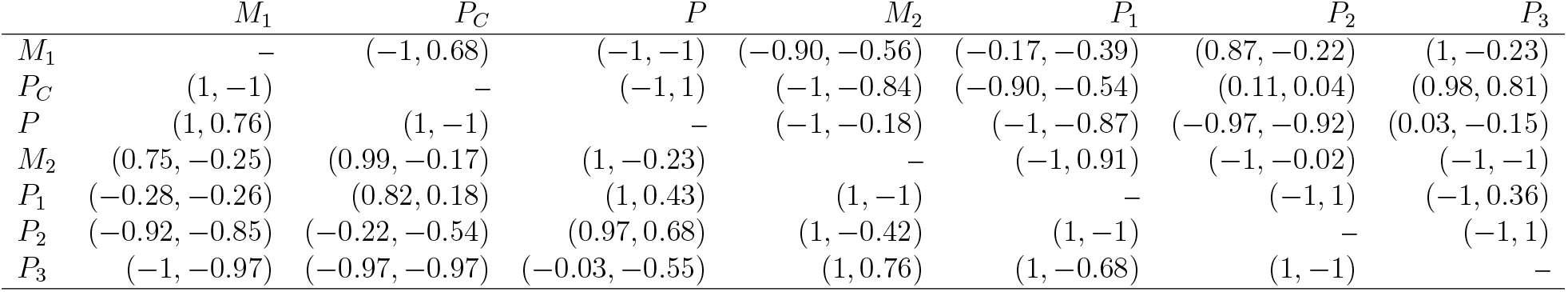
Calculated regulation-detection scores for the Kim-Forger, Goodwin independent cycle example (Fig. 2E).

| | $M_1$ | $P_C$ | $P$ | $M_2$ | $P_1$ | $P_2$ | $P_3$ |
| --- | --- | --- | --- | --- | --- | --- | --- |
| $M_1$ | – | (–1, 0.68) | (–1, –1) | (–0.90, –0.56) | (–0.17, –0.39) | (0.87, –0.22) | (1, –0.23) |
| $P_C$ | (1, –1) | – | (–1, 1) | (–1, –0.84) | (–0.90, –0.54) | (0.11, 0.04) | (0.98, 0.81) |
| $P$ | (1, 0.76) | (1, –1) | – | (–1, –0.18) | (–1, –0.87) | (–0.97, –0.92) | (0.03, –0.15) |
| $M_2$ | (0.75, –0.25) | (0.99, –0.17) | (1, –0.23) | – | (–1, 0.91) | (–1, –0.02) | (–1, –1) |
| $P_1$ | (–0.28, –0.26) | (0.82, 0.18) | (1, 0.43) | (1, –1) | – | (–1, 1) | (–1, 0.36) |
| $P_2$ | (–0.92, –0.85) | (–0.22, –0.54) | (0.97, 0.68) | (1, –0.42) | (1, –1) | – | (–1, 1) |
| $P_3$ | (–1, –0.97) | (–0.97, –0.97) | (–0.03, –0.55) | (1, 0.76) | (1, –0.68) | (1, –1) | – |

**Table S4:**
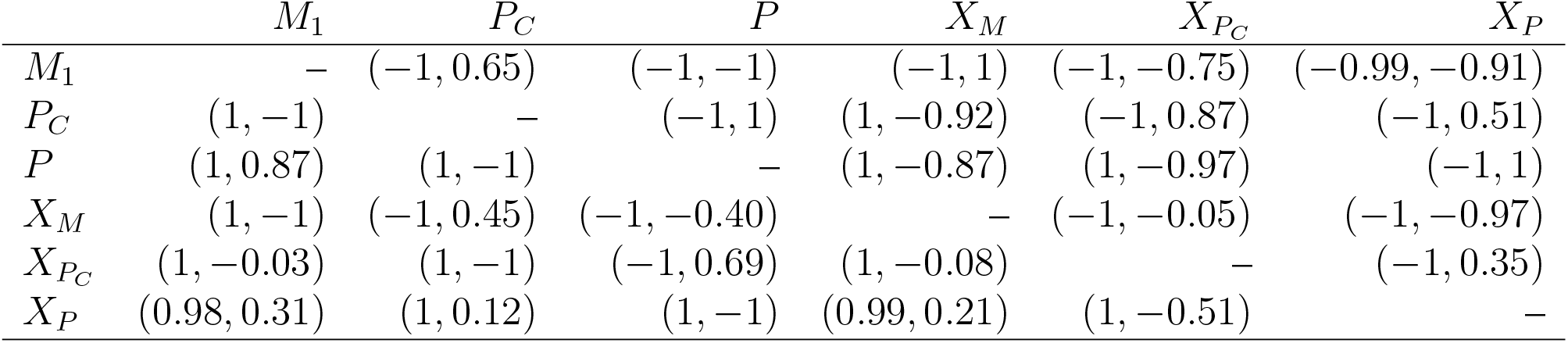
Calculated regulation-detection scores for the Kim-Forger with outputs example (Fig. 2F).

| | $M_1$ | $P_C$ | $P$ | $X_M$ | $X_{P_C}$ | $X_P$ |
| --- | --- | --- | --- | --- | --- | --- |
| $M_1$ | – | (–1, 0.65) | (–1, –1) | (–1, 1) | (–1, –0.75) | (–0.99, –0.91) |
| $P_C$ | (1, –1) | – | (–1, 1) | (1, –0.92) | (–1, 0.87) | (–1, 0.51) |
| $P$ | (1, 0.87) | (1, –1) | – | (1, –0.87) | (1, –0.97) | (–1, 1) |
| $X_M$ | (1, –1) | (–1, 0.45) | (–1, –0.40) | – | (–1, –0.05) | (–1, –0.97) |
| $X_{P_C}$ | (1, –0.03) | (1, –1) | (–1, 0.69) | (1, –0.08) | – | (–1, 0.35) |
| $X_P$ | (0.98, 0.31) | (1, 0.12) | (1, –1) | (0.99, 0.21) | (1, –0.51) | – |

**Table S5:**
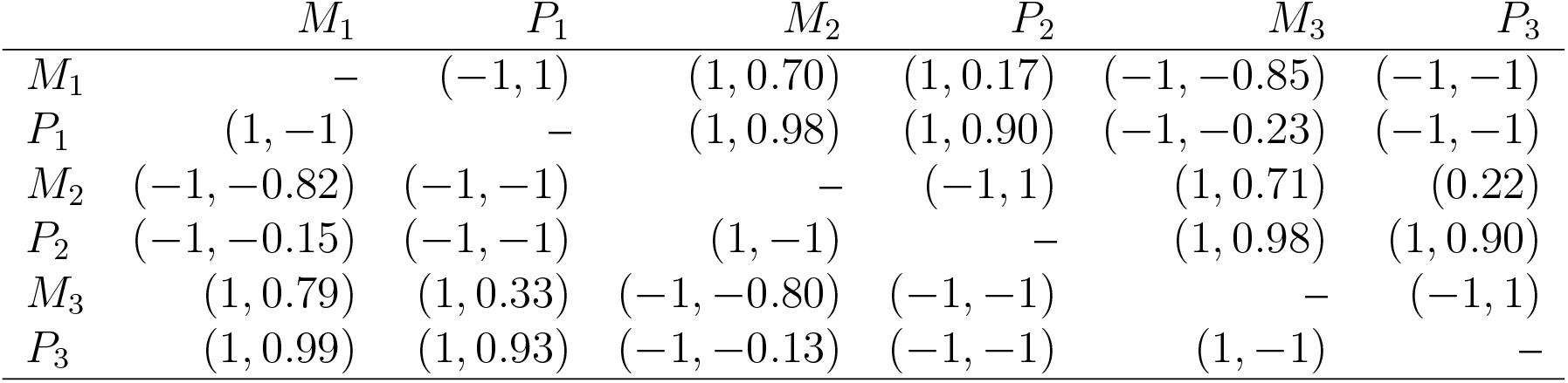
Calculated regulation-detection scores for the repressilator example (Fig. 2G).

| | $M_1$ | $P_1$ | $M_2$ | $P_2$ | $M_3$ | $P_3$ |
| --- | --- | --- | --- | --- | --- | --- |
| $M_1$ | – | (–1, 1) | (1, 0.70) | (1, 0.17) | (–1, –0.85) | (–1, –1) |
| $P_1$ | (1, –1) | – | (1, 0.98) | (1, 0.90) | (–1, –0.23) | (–1, –1) |
| $M_2$ | (–1, –0.82) | (–1, –1) | – | (–1, 1) | (1, 0.71) | (0.22) |
| $P_2$ | (–1, –0.15) | (–1, –1) | (1, –1) | – | (1, 0.98) | (1, 0.90) |
| $M_3$ | (1, 0.79) | (1, 0.33) | (–1, –0.80) | (–1, –1) | – | (–1, 1) |
| $P_3$ | (1, 0.99) | (1, 0.93) | (–1, –0.13) | (–1, –1) | (1, –1) | – |

**Table S6:** Calculated regulation-detection scores for the experimental repressilator example (Fig. 4B).

| | $P_1$ | $P_2$ | $P_3$ |
| --- | --- | --- | --- |
| $P_1$ | – | $(-0.99, -0.92)$ | $(0.21, -0.006)$ |
| $P_2$ | $(0.75, 0.71)$ | – | $(-1, -0.98)$ |
| $P_3$ | $(-1, -0.93)$ | $(0.98, -0.04)$ | – |

**Table S7:** Calculated regulation-detection scores for the mixed repressilator example (Fig. 4C).

| | $P_1^1$ | $P_2^1$ | $P_3^1$ | $P_1^2$ | $P_2^2$ | $P_3^2$ |
| --- | --- | --- | --- | --- | --- | --- |
| $P_1^1$ | – | $(-1, -0.96)$ | $(0.56, -0.18)$ | $(-0.83, -0.01)$ | $(0.97, -0.57)$ | $(-1, 0.37)$ |
| $P_2^1$ | $(0.83, 0.88)$ | – | $(-1, -0.97)$ | $(-0.65, 0.99)$ | $(-0.19, 0.18)$ | $(0.96, 0.14)$ |
| $P_3^1$ | $(-1, -1)$ | $(0.96, 0.02)$ | – | $(1, -0.03)$ | $(-1, 0.16)$ | $(-0.96, 0.05)$ |
| $P_1^2$ | $(-0.69, 0.37)$ | $(0.93, -0.74)$ | $(-1, -0.24)$ | – | $(-0.99, -0.91)$ | $(-0.04, -0.09)$ |
| $P_2^2$ | $(-0.97, 0.98)$ | $(0.15, 0.38)$ | $(1, -0.65)$ | $(0.76, 0.61)$ | – | $(-1, -0.96)$ |
| $P_3^2$ | $(1, -0.31)$ | $(-1, -0.11)$ | $(-0.67, 0.23)$ | $(-1, -0.91)$ | $(0.97, -0.04)$ | – |

**Table S8:**
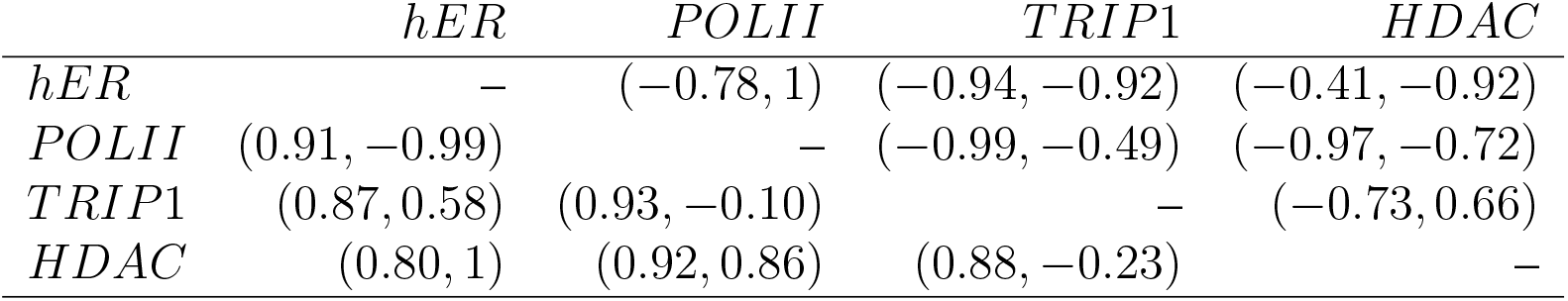
Calculated regulation-detection scores for the estradiol example (Fig. 4D).

| | $hER$ | $POLII$ | $TRIP1$ | $HDAC$ |
| --- | --- | --- | --- | --- |
| $hER$ | – | $(-0.78, 1)$ | $(-0.94, -0.92)$ | $(-0.41, -0.92)$ |
| $POLII$ | $(0.91, -0.99)$ | – | $(-0.99, -0.49)$ | $(-0.97, -0.72)$ |
| $TRIP1$ | $(0.87, 0.58)$ | $(0.93, -0.10)$ | – | $(-0.73, 0.66)$ |
| $HDAC$ | $(0.80, 1)$ | $(0.92, 0.86)$ | $(0.88, -0.23)$ | – |

